# StruCloze: A Unified Framework for Backmapping and Inpainting of Biomolecules

**DOI:** 10.1101/2025.06.26.661889

**Authors:** Junjie Zhu, Zirui Fan, Zhengxin Li, Zhuoqi Zheng, Kresten Lindorff-Larsen, Hai-Feng Chen

## Abstract

Atomistic resolution is essential for understanding biomolecular structure and function, yet coarse-grained (CG) models remain indispensable for simulating large and dynamic systems. Reconstructing accurate all-atom structures from CG representations, particularly across varied CG schemes and biomolecular types, remains a fundamental challenge. Moreover, flexible or disordered regions may well suffer from failure in structure modeling, making inpainting missing regions another challenging task. Here, we present StruCloze, a deep learning framework for reconstructing atomistic structures from CG models and inpainting missing regions for both proteins and nucleic acids. StruCloze generalizes across various CG levels and biomolecule types on single pretraining, with minimal fine-tuning required for optimal performance on specific representations. It achieves state-of-the-art accuracy in reconstructing both protein and nucleic acid structures, demonstrates superior transferability and speed compared to existing methods. Leveraging masked learning strategy, StruCloze also excels at inpainting structurally missing regions in structures, offering a practical tool for structural refinement and integrative modeling. Our framework provides a general solution for bridging reduced or incomplete representations with full atomistic detail of biomolecular structures, enabling rapid local structure prediction and further analysis on system dynamics.

## Introduction

Biomolecular interactions underlie the majority of cellular functions, enabling precise and rapid cellular responses to internal and external stimuli^1,2^. Understanding the structural details of biomolecular complexes is crucial not only for deciphering biological mechanisms but also for the rational design of targeted therapeutics^3,4^. While experimental techniques such as X-ray crystallography and cryogenic electron microscopy (cryo-EM) have greatly advanced structural biology, they remain resource-intensive and struggle to capture flexible elements like intrinsically disordered regions (IDRs) and RNAs^5–7^.

To complement experimental efforts, computational modeling and molecular simulations have become indispensable for exploring biomolecular dynamics, interactions and mechanisms^8–10^. All-atom simulations provide detailed insights but suffer from high computational complexity for large systems. Despite major breakthroughs such as AlphaFold2 and AlphaFold3, current structure prediction methods remain largely limited to static representations and cannot fully account for conformational diversity or dynamic assembly processes^11,12^.

Coarse-grained (CG) modeling provides an efficient alternative by reducing structural complexity, thereby facilitating simulations of large-scale biomolecular behavior^13^. CG representations also better accommodate flexible and low-resolution regions, making them particularly useful for integrative modeling with experimental data^14^. Typically, CG models range from ultra-coarse models (one bead per biomolecule^15^) to more detailed ones (residue-based models^16–19^ and multiple beads per residue^20–22)^, with trade-offs between accuracy, transferability and speed.

However, coarse-graining inevitably sacrifices atomic detail, making reconstruction of all-atom structures from CG models a critical step for downstream analysis. While hydrogen atoms can often be inferred from local geometries, accurate placement of remaining heavy atoms remains a central challenge^23^. For high-resolution protein CG models, geometry-based reconstruction rules can generally recover all-atom structures with reasonable accuracy^24^. However, the task becomes significantly more complex for coarser representations such as residue-based models. Methods like PULCHRA typically reconstruct the protein backbone from CA traces first, followed by the prediction of side-chain orientations^25^. Given the higher flexibility of protein side chains, an additional set of studies have focused specifically on reconstructing side-chain structures or generating realistic side-chain conformational ensembles^26,27^. For nucleic acids, existing methods such as ARENA and ABC2A typically rely on template matching followed by iterative optimization to reconstruct all-atom structures^28,29^. In most cases, extensive structural optimization is required to avoid clashes and produce more confident all-atom models.

Recent machine learning efforts—including models like CGVAE, GenZProt, and cg2all—have demonstrated high accuracy in reconstructing protein structures from coarse-grained inputs^30–32^. Nevertheless, these models often lack generalizability: they are typically trained on one specific CG representation and require extensive retraining when applied to different CG schemes. Moreover, accurate atomistic reconstruction for nucleic acids with machine learning remains an underexplored area.

In this work, we introduce StruCloze, a transferable deep learning framework for reconstructing all-atom structures of both proteins and nucleic acids from various CG representations. Through a single pretraining process, StruCloze learns generalizable mappings across CG levels, while simple fine-tuning enables further performance gains on specific CG schemes. StruCloze outperforms state-of-the-art methods in reconstructing structures for both proteins and nucleic acids. Beyond reconstruction, StruCloze is capable of inferring missing regions in low-resolution experimental structures, offering a unified solution for structural refinement and restoration.

## Results

### Overview

We introduce StruCloze, a general framework for reconstructing all-atom biomolecular structures from incomplete or coarse-grained inputs (Figure 1A). StruCloze is designed to accommodate a wide range of input CG representations, including coarse-grained models with one bead per residue (e.g., Cα for proteins, C4′ for RNAs, and center-of-mass representations) as well as more detailed multi-bead models (e.g., CALVADOS-RNA, oxDNA, MARTINI). In addition to coarse representations, StruCloze can effectively infer the positions of unresolved regions, such as missing residues or side chains.

**Figure 1.**
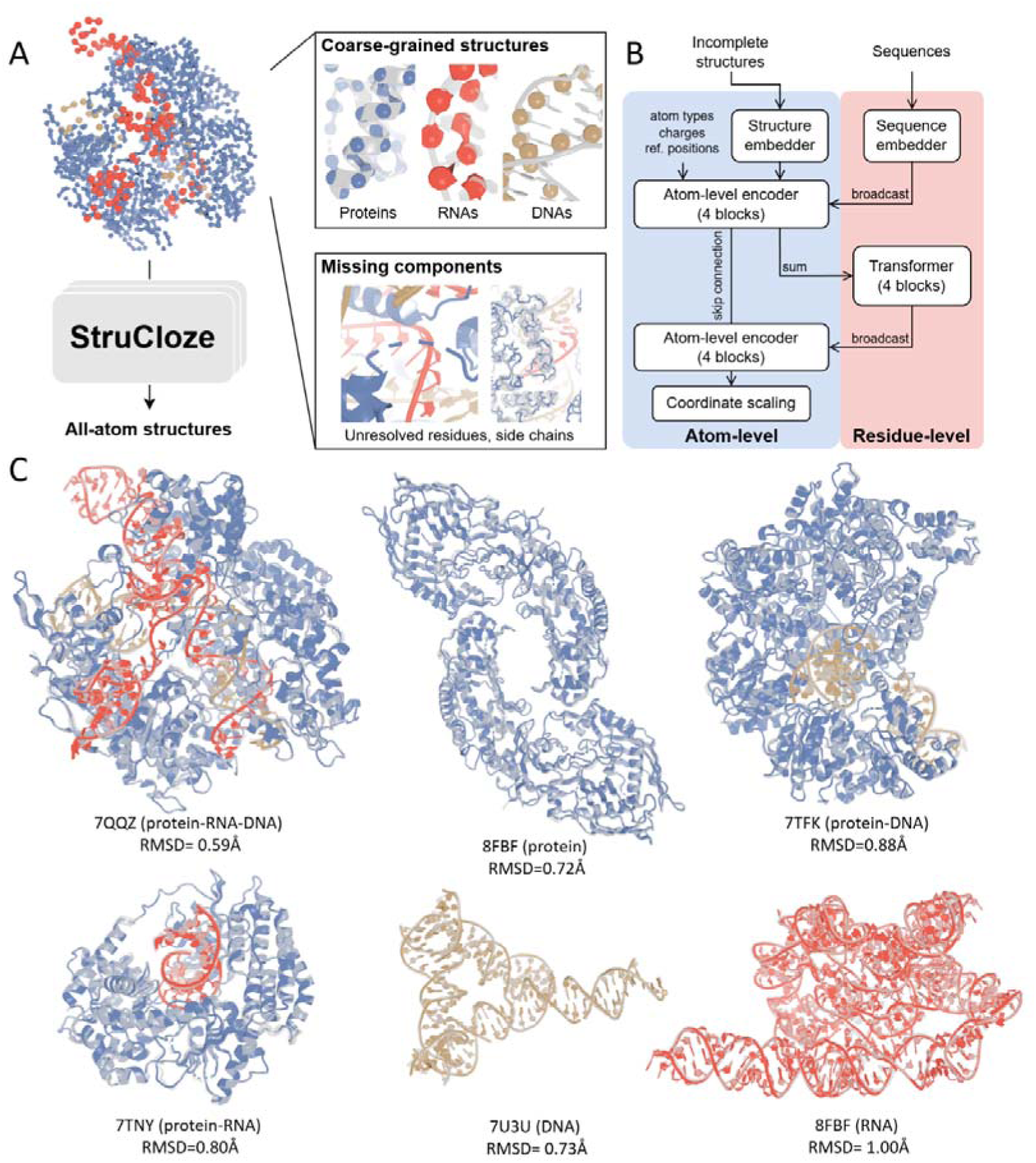
Overview of StruCloze. (A) StruCloze addresses various forms of incomplete biomolecular structures. (B) Schematic representation of the StruCloze architecture. (C) Examples of full atomic reconstructions generated by StruCloze, compared with experimentally resolved structures (grey).

The architecture of StruCloze adopts a dual-level design (Figure 1B). The atom-level module processes structural features and spatial coordinates, using stacked encoder blocks to extract local atomic context. In parallel, the residue-level module captures global sequence-level interactions via a Transformer-based encoder. Cross-level broadcasting enables joint reasoning across both atom and residue levels, while skip connections and coordinate scaling ensure stability and precision in structure prediction.

StruCloze was initially pretrained on center-of-mass (COM) representations, which offer a simple yet informative scheme applicable to both proteins and nucleic acids. We displayed several reconstructed biomolecular assemblies from COM inputs (Figure 1C). The reconstructed all-atom structures showed high accuracy, with RMSD values typically below 1 Å relative to experimental references. Implementation details and training procedures are provided in Materials and Methods.

### Performance on Cα/C4’ and COM

To systematically evaluate StruCloze and compare it with prior backmapping methods, we curated a test set of 8,066 biomolecular structures from the Protein Data Bank (PDB). The curation procedure follows that of AlphaFold3^11^, with ligands ignored during processing. We first assessed the performance of StruCloze on commonly used single-particle-per-residue CG representations, Cα/C4′ and center-of-mass (COM), and benchmarked it against the current state-of-the-art method on protein structures, cg2all. cg2all represents protein backbones as rigid bodies and reconstructs them via predicted rigid transformations, while side chains are rebuilt through predicted torsion angles. This scheme is effective for proteins but struggles to generalize to nucleotides, which exhibit higher internal flexibility.

The test set was categorized into protein-only (7,310 structures), nucleic acid-only (78 structures), and protein–nucleic acid hybrid complexes (678 structures). We computed RMSD values for all heavy atoms reconstructed by both StruCloze and cg2all on protein structures. StruCloze achieved a comparable reconstruction RMSD for the Cα representation compared to cg2all (1.03 Å vs. 1.06 Å) but yielded more steric clashes (0.24% vs. 0.15%, Figure 2A). For the COM representation, StruCloze outperformed cg2all on both RMSD and clash ratio, and both methods showed improved accuracy over the Cα-based reconstructions (Figure 2B, 2D). This improvement is likely due to the COM representation implicitly capturing side-chain atomic positions, providing richer geometric cues^32^. In contrast, Cα-based representations contain less side-chain information^33^, leading to greater uncertainty in modeling side-chain conformations. This observation aligns with prior coarse-grained modeling studies suggesting that COM representations better preserve side-chain interactions and stabilize structured regions^34^.

**Figure 2.**
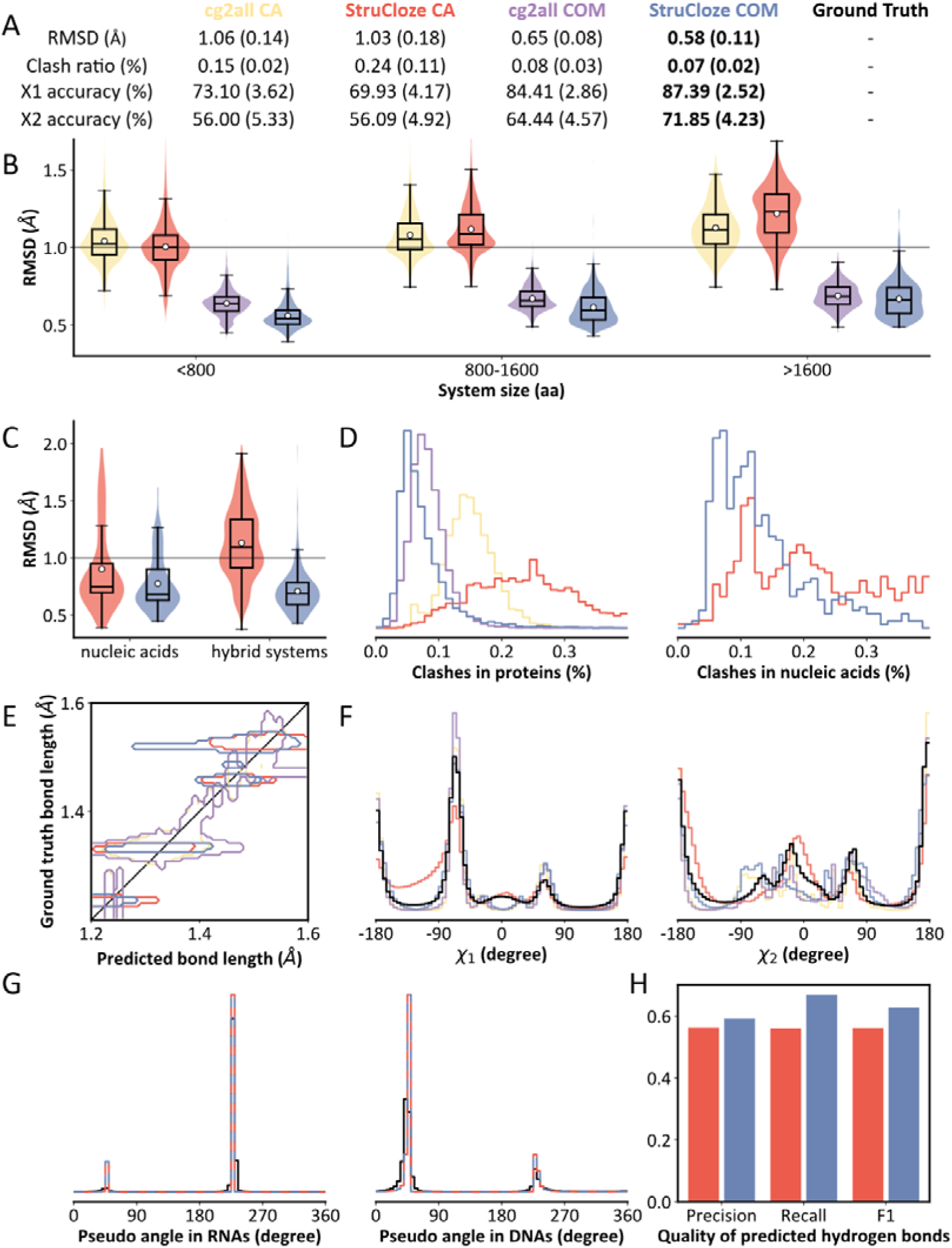
Performance of StruCloze for Cα/C4’ and COM representations. (A) Summary of RMSD, clash ratio and, accuracy on proteins. Standard deviations are given in the parentheses. (B) RMSD distributions on proteins. (C) RMSD distributions on nucleic acids and hybrid systems. (D) Clash ratio distributions on proteins and nucleic acids. (E) Bond lengths. (F) Protein side chain dihedral *χ*_1_ and *χ*_2_. (G) Pseudo angles of ribose in nucleic acids, representing C2’ or C3’-endo conformations. (H) Quality of predicted hydrogen bonds.

In nucleic acid and hybrid systems, StruCloze maintained high accuracy, with RMSD values from both C4′ and COM reconstructions only slightly higher than those observed for proteins, and all reconstructions remained below 2 Å RMSD (Figure 2C). Clash ratios in these systems were also comparable to those in protein-only cases, indicating that the model—trained directly on atomic coordinates and supplemented with nucleic acid training data—successfully generalizes from proteins to nucleic acids (Figure 2D). In contrast, cg2all’s rigid-body framework and other internal-coordinate-based methods like GenZProt exhibit limited adaptability, particularly for modeling the flexible backbone of nucleic acids. StruCloze thus demonstrates unprecedented transferability and robustness across biomolecules.

In evaluating clash ratios, we only considered atom pairs at least 3 Å apart in the ground truth structures, excluding bonded atoms. To further assess geometric fidelity, we analyzed predicted distances between bonded atom pairs in proteins (1.2–1.6 Å range, Figure 2E, Figure S1). StruCloze accurately reproduced typical bond lengths along the backbone (N–Cα ∼1.46 Å, Cα–C ∼1.53 Å, peptide bond ∼1.33 Å), closely matching cg2all. However, for side-chain C–C bonds, we observed a mild underestimation of interatomic distances. Since StruCloze is supervised directly on atomic coordinates, it lacks explicit constraints on bond lengths or torsions, leading to suboptimal optimization of local geometry. What’s worth noting is that the test structures were derived from high-resolution crystallographic models, which tend to exhibit idealized bond lengths. In contrast, physiological or simulated conformations typically display broader bond-length distributions^35^.

Beyond bond lengths, torsional distributions provide another critical indicator of structural plausibility. We analyzed backbone dihedral angles (□, ψ) and all side-chain torsions (χ1–χ4) in both ground truth and reconstructed structures. StruCloze faithfully reproduced backbone dihedral distributions, with sampling well-concentrated in favored regions (Figure S2). Predictions for χ1 and χ2 were also highly accurate (Figure 2F). However, for χ3 and χ4 in long, flexible side chains such as Arg and Lys, the model failed to capture certain regions in the upper-left and lower-right quadrants of the dihedral space (Figure S3). This is likely due to the sparse representation of these distal torsions in the CG input and their low frequency in the training data, particularly for χ4. Modeling these distant torsions remains challenging for deep learning approaches. One should also note that χ2–χ4 torsions are less well-defined due to their high conformational flexibility and are therefore not determined uniquely by the backbone structure.

In nucleic acids, we similarly evaluated both backbone and base modeling accuracy. For the backbone, we examined the virtual torsion angle that defines ribose puckering, distinguishing between C2′-endo and C3′-endo conformations. StruCloze accurately modeled the distinct ribose geometries in DNA and RNA from both C4’ and COM representation (Figure 2G). For base pairing, we assessed the model’s ability to predict hydrogen-bonding atom pairs using binary classification metrics. StruCloze achieved high accuracy in recovering native base-pairing interactions, demonstrating its potential to reconstruct functional nucleic acid conformations with atomic fidelity (Figure 2H).

### Direct prediction and fine-tuning across CG representations

We further evaluated the transferability of StruCloze to alternative CG representations without additional fine-tuning. For multi-bead-per-residue models, CG bead coordinates were computed for both the all-atom structures and the corresponding CCD reference structures. The references were aligned to the target CG configurations using the Kabsch algorithm, and were used as input to the model.

We first assessed performance on nucleic acid systems, selecting three widely used RNA CG representations: IsRNA1, IsRNA2, and CALVADOS-RNA. Due to the limited availability of RNA test systems, we also applied these representations to DNA, evaluating model performance across 78 systems. The first two CG schemes—primarily used for RNA structure prediction—represent each nucleotide with 2 beads for backbone and 2–3 beads for bases. Given the rich CG information, transferring StruCloze directly to IsRNA1 led to no significant increase in reconstruction error (0.81□Å on IsRNA1 vs. 0.77□Å on COM; *p* = 0.21). A similar trend was observed for IsRNA2 (0.82□Å; *p* = 0.14). However, for CALVADOS-RNA, which is more coarsely grained with only 2 beads per nucleotide, direct transfer resulted in a significant accuracy drop (1.11□Å; *p* < 0.01), likely due to large discrepancies in residue position initialization between CALVADOS-RNA and COM formats (Figure 3A). The clash ratio showed consistent trends with RMSD across representations (Figure 3B).

**Figure 3.**
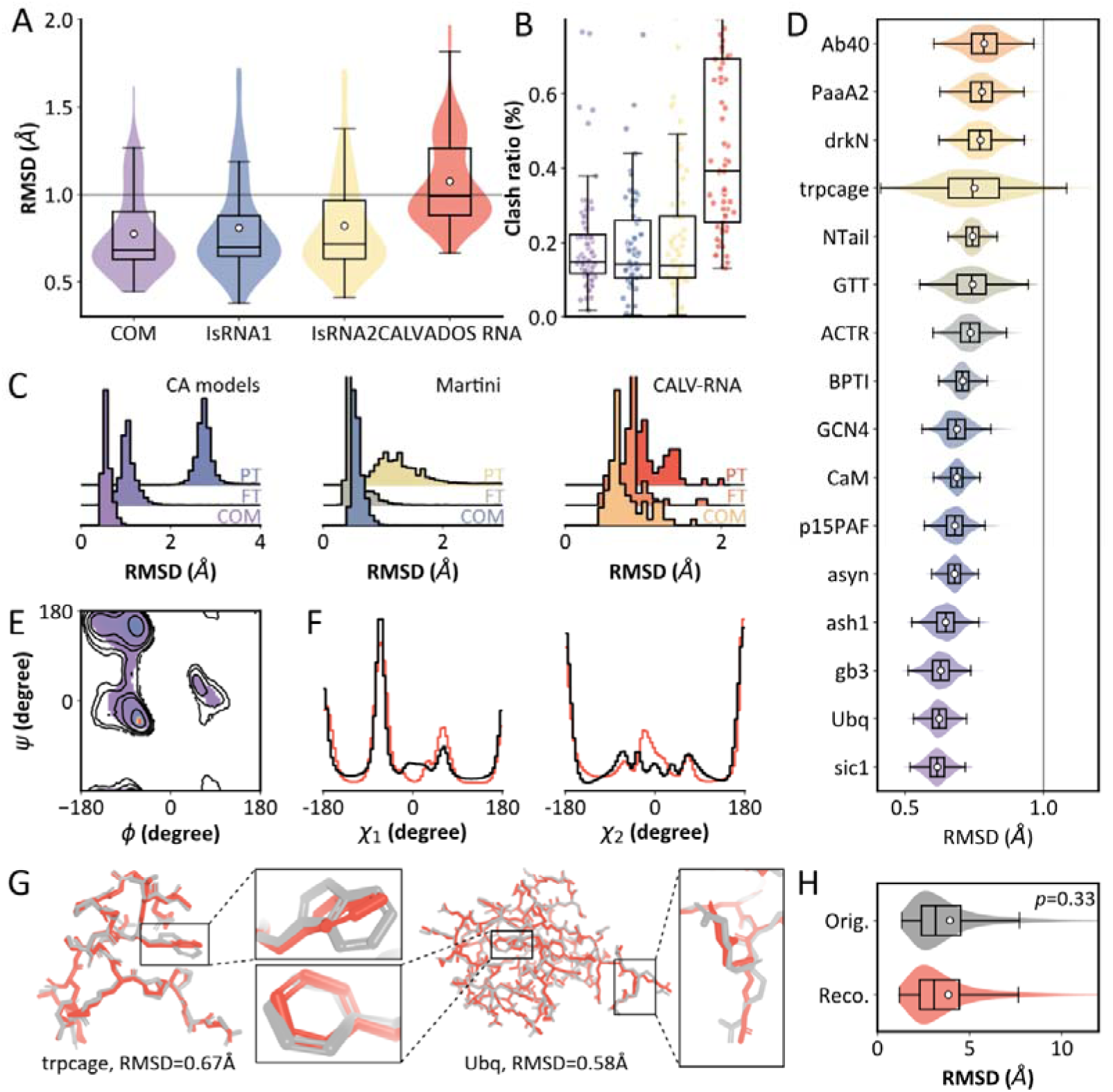
Performance of StruCloze across CG models and MD trajectories. (A) Reconstruction RMSD from direct predictions on RNA CG models with multiple beads per residue. (B) Clash ratio for reconstruction from RNA CG models. (C) Effect of finetuning on reconstruction performance. PT (top) denotes direct prediction of pretrained model, FT (middle) denotes finetuned model, and COM (bottom) shows pretrained performance on the COM representation as a reference. (D) Reconstruction RMSD on 16 IDP trajectories. (E) Ramachandran plot of reconstructed IDP trajectories. Distribution of the original trajectories is shown as contours. (F) Distributions of χ angles in original (black) and reconstructed (red) IDP trajectories. (G) Representative prediction errors made by StruCloze. Predicted structures are shown in red, and ground truth structures in grey. (H) Pairwise RMSD within trajectories of original and reconstructed trajectories from ATLAS dataset.

In protein systems, transferring the COM-trained model to Cα and Martini models also led to increased errors, with a more pronounced drop in accuracy. A likely reason is the training data being protein-dominated, which resulted in more optimization iterations for protein systems and thus more severe overfitting to the COM representation, limiting generalization to other CG formats.

Despite the elevated error in certain CG formats under direct transfer, a short fine-tuning step on the target representation (40 epochs versus 200 during pretraining) is sufficient to effectively adapt StruCloze. As shown in Figure 3C, post-finetuning reconstruction RMSDs were significantly reduced for CA, Martini, and CALVADOS-RNA. Notably, owing to Martini models’ richer bead-level detail, the fine-tuned model outperformed the original COM model. These results highlight the strong transferability of StruCloze and its capacity to rapidly and accurately reconstruct all-atom structures from a diverse range of CG inputs.

### Prediction for MD trajectories

Practical applications of backmapping methods often involve reconstructing conformational ensembles derived from molecular dynamics (MD) simulations. We further assessed StruCloze on all-atom MD trajectories generated with a99SB-disp^36^. Twenty protein systems were selected, and 1,000 frames were sampled at even intervals for evaluation. For each sampled frame, we reconstructed the all-atom structure from the corresponding COM trace using StruCloze and computed the RMSD with respect to the ground-truth atomistic conformation.

Across all systems, reconstruction RMSDs remained below 1□Å for the majority of frames, indicating good overall accuracy (Figure 3D). However, a slight increase in reconstruction error was observed compared to results on crystal structures, suggesting that the model trained exclusively on static crystal structures may well be inherently biased toward low-energy, canonical conformations, and do not fully capture the broader conformational diversity sampled during MD simulations.

Torsional angle distributions are further analyzed for both MD trajectories and corresponding reconstructed structures (Figure 3E, 3F). MD trajectories exhibited broader distributions and presented more rotamer outliers resulting from thermal fluctuations. In contrast, torsional distributions of StruCloze-reconstructed structures were markedly more constrained, closely resembling those found in crystallographic ensembles. This indicates a tendency of the model toward low-energy rotamers, thus failing to recover the dynamics inherent in MD trajectories.

Further analysis revealed specific failure modes (Figure 3G, Figure S4). A disproportionate number of reconstruction errors were localized at phenylalanine and tryptophan, both of which contain planar aromatic side chains. Side-chain orientations for these residues were more frequently incorrect compared to other residue types in reconstructed trajectories. Moreover, residues at chain termini were more error-prone. Reduced local context and large Cartesian coordinate values result in larger deviations in both backbone and side-chain geometry in these terminal residues.

While reconstructing a structure within a specific RMSD is important, it’s equally crucial that the reconstructed structure is closer to its original counterpart than to other structures within the same simulation. Therefore, we further analyze the differences between reconstructed and original trajectories on 1,390 structured protein trajectories from the ATLAS dataset^37^. For each system, 30 frames were extracted at consistent intervals. Structured proteins, being more stable in simulation than IDPs, offer a more sensitive test for errors in reconstructing pairwise similarity among frames. Analysis of pairwise RMSD within trajectories showed no significant difference between reconstructed and original trajectories (*p* = 0.33, Figure 3H). This suggests that StruCloze effectively preserves the intricate pairwise relationships between correlated conformations.

### Sampling atomistic structures for condensates

We further focused on two intrinsically disordered region (IDR) systems: one from leucine-rich repeat-containing protein 37A2 (residues 1082–1700 from UniProt entry A6NM11, referred to as 37A2), and one from WD repeat and FYVE domain-containing protein 3 (residues 1899–1953 from UniProt entry Q8IZQ1, referred to as FYVE). 37A2 has been shown to exhibit strong phase separation propensity (ΔG = -8.44*k_B_*T), while FYVE tends to remain in the dilute phase (ΔG > 0*k_B_*T). CG trajectories simulated using CALVADOS2 with Cα representations were obtained from previous work by Sören von Bülow et al^38^.

A direct observation from the backmapped trajectories is the significantly higher number of inter-chain contacts in the 37A2 system compared to FYVE (Figure 4A, B), consistent with the tendency of 37A2 to form a dense phase, whereas FYVE remains in the dilute phase. Visualizations of the reconstructed structures also reflect this phase behavior. It’s essential to note that this global phase separation tendency is primarily determined by the CG simulation itself, StruCloze introduces little substantial errors in residue placement.

**Figure 4.**
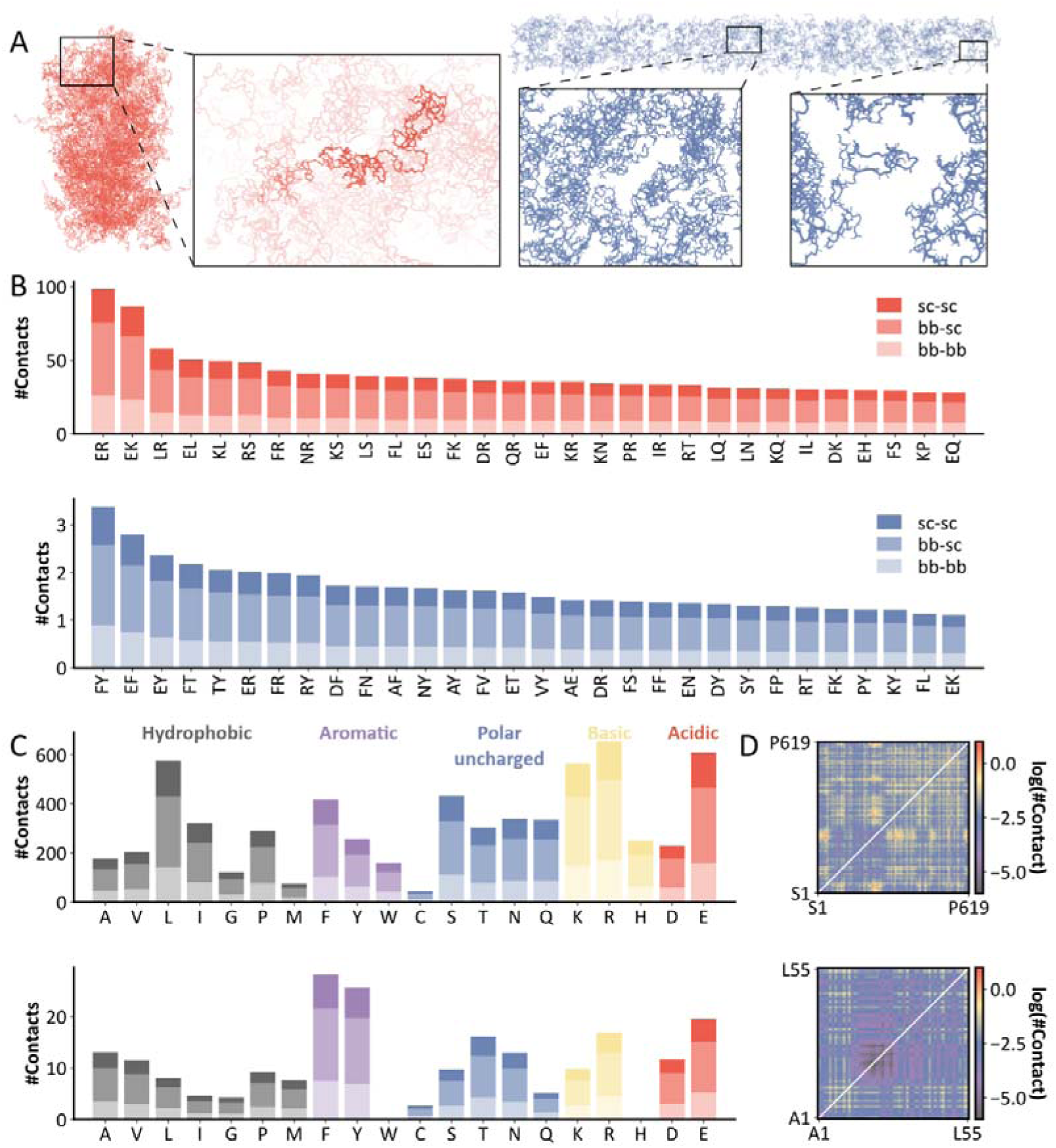
Analysis on reconstructed all-atom trajectories for two condensates. (A) Structure of condensate formed by 37A2 (red) and FYVE (blue) in the dilute phase. (B) Top 30 inter-chain contact types in 37A2 (red) and FYVE (blue). (C) Number of inter-chain contacts contributed by 20 amino acids in 37A2 (top) and FYVE (bottom). (D) Average inter-chain contact in 37A2 (top) and FYVE (bottom).

We further analyzed the interaction types in the reconstructed all-atom structures (Figure 4C). In 37A2, the dominant interacting residue pairs were E–R and E–K, representing classical charge–charge attractions. In contrast to these electrostatic interactions, aromatic-mediated contacts were far less frequent. These findings suggest that the phase separation observed in 37A2 is primarily driven by electrostatic interactions. On the other hand, FYVE exhibited very few electrostatic contacts. Instead, interactions were primarily of the aromatic–aromatic (F–Y) and aromatic–cationic (E–F, E–Y) types. The number of inter-chain contacts in FYVE was minimal, and no obvious dense phase was observed.

We further examined the average contact maps (Figure 4D). In 37A2, the region spanning P216–M256 contributed most of the interchain interactions, while other regions showed relatively sparse contacts, suggesting that this segment may serve as a key driver for condensate stability. In FYVE, the interactions were more evenly distributed. The N-terminal residues F5, F7, R10, and Y12 were involved in relatively frequent interactions, although still at much lower levels than observed in 37A2.

Additionally, we performed backmapping on a hybrid system involving proteins (FUSRGG3) and disordered RNAs (polyU40)^22^. While the protein components were reconstructed accurately, the RNA structures posed challenges (Figure S5). Specifically, StruCloze tended to generate nucleotide conformations resembling those of canonical double-stranded RNA. However, the CG representations used in the simulation reflected highly flexible, disordered RNA structures whose nucleotide conformations deviate substantially from crystal-like geometries. As a result, although the model could roughly place each nucleotide, it failed to accurately restore the ribose geometry and phosphate backbone connectivity, primarily due to errors in sugar positioning and puckering. Modeling such highly flexible RNA structures will require further finetuning with additional dynamic training data to capture their structural heterogeneity more effectively.

### Predicting missing components for incomplete experimental structures

During evaluation, we observed that prior deep learning methods often suffer from limited generalizability. For example, internal-coordinate-based models such as GenZProt may fail to reconstruct structures even within the same CG representation when the input deviates significantly from training data, indicating poor robustness^31^. cg2all trains separate models for different common protein CG schemes and achieves high accuracy within each, but lacks transferability across representations^32^. This suggests that many current backmapping networks are overfitted to specific CG formats. A similar issue was also observed in StruCloze. When residues are missing in the input structure, the model tends to place them at positions corresponding to the default mask values (typically zero), reflecting strong dependence on the provided CG coordinates.

Nevertheless, we argue that the modeling framework of StruCloze should, in principle, be transferable and capable of inferring missing residues from surrounding context. To this end, we applied a masked self-supervision strategy, randomly masking 10% of residues during finetuning and supervising model prediction on these positions. To evaluate the performance of StruCloze in reconstructing unresolved regions, we first applied random masking at various levels to test structures and measured reconstruction RMSD on all heavy atoms. When using the finetuned model, RMSD remained below 1DÅ at mask ratios ≤30%. Although the reconstruction quality slightly decreased as the mask ratio increased, RMSD values rarely exceeded 2DÅ even when 50% residues were masked, and the clash ratio remained consistently low (Figure 5A). Since masked regions are assigned zero coordinates as input, the pretrained model tends to predict these regions near the origin as it treats the masks the same as CG representations. In contrast, the finetuned model more accurately restores the missing residues to their correct positions (Figure 4B).

**Figure 5.**
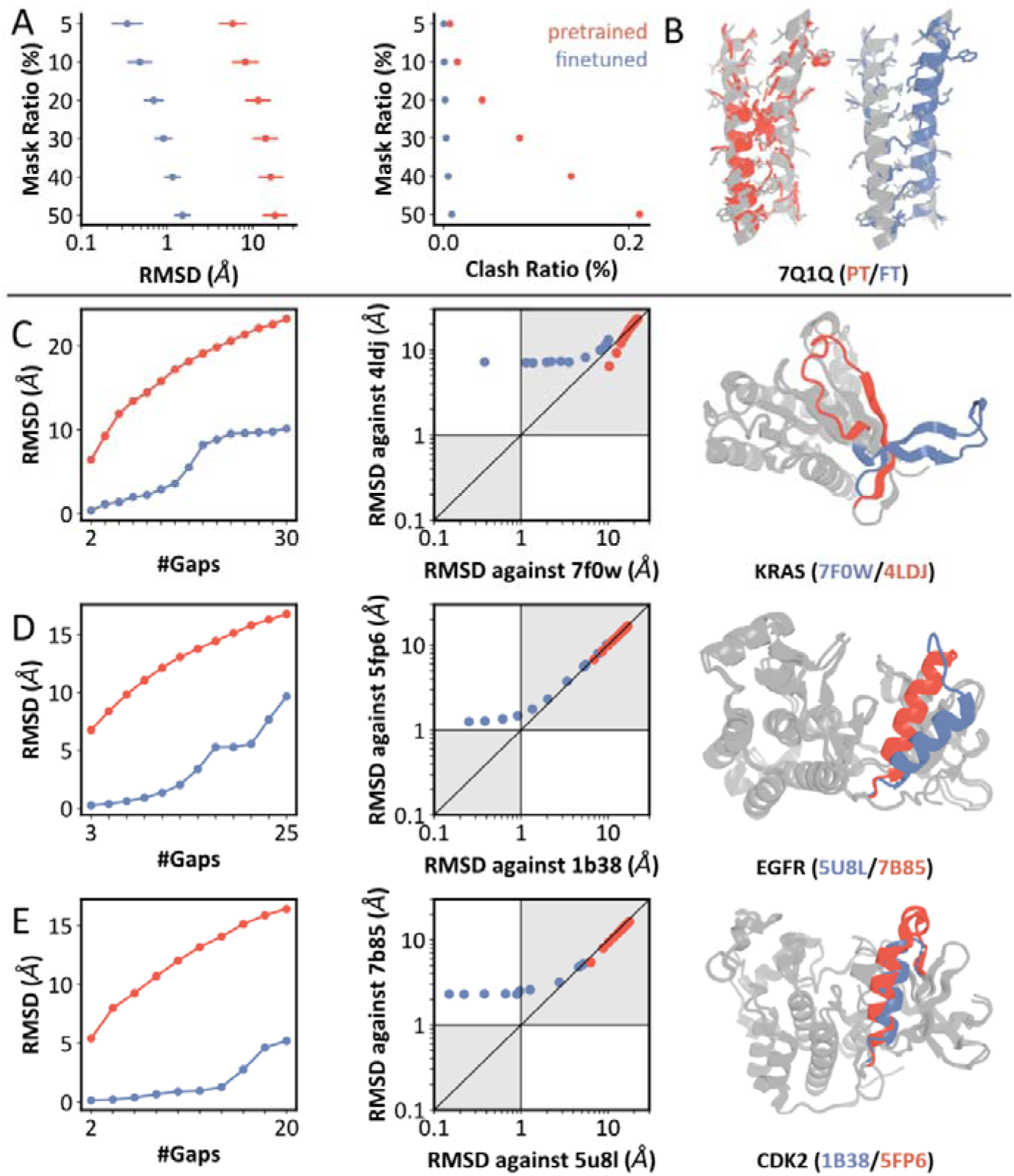
Performance of StruCloze in predicting unresolved regions. (A) RMSD and clash ratio at different mask ratios. (B) Predicted structures for 7Q1Q at 10% mask ratio using pretrained and finetuned models. (C-E) RMSD for (C) KRAS, (D) EGFR and (E) CDK2 at different gap lengths. Left: RMSD to the original structure. Middle: RMSD to both reference conformations. Right: Reference structures. Red and blue scatters denote inpainting from reference structure of certain color.

To further explore the ability and limitations of StruCloze in inpainting missing regions, we designed tests on three allosteric protein systems: KRAS, EGFR, and CDK2. For each protein, we selected two distinct reference conformations, none of which appeared in the training set. These conformations are globally similar but contain a local difference in a continuous segment of ∼30 residues. We masked segments of varying lengths from one conformation and evaluated the model’s ability to reconstruct the missing region (Figures 5C–E). Using RMSD□≤□1□Å as the threshold for accurate reconstruction, the model successfully predicted masked segments of length 2, 9, and 10 residues in KRAS, EGFR, and CDK2, respectively—each corresponding only to one of the two reference structures. For longer masked regions, the model still tended to predict the missing residues near the origin, leading to large RMSD values against both reference structures.

On one hand, the lack of reconstruction toward alternative conformations as the masked region length increased suggests that StruCloze behaves deterministically and lacks generative diversity in its predictions. On the other hand, we observed that inpainting from different conformations could lead to reconstruction failure, even though the pretrained model successfully reconstructs both conformations from coarse-grained input. Previous studies have shown that AlphaFold2 can predict alternative conformations of the same protein due to its memorization of certain structural patterns during training^39^. A similar mechanism is supposed to work for StruCloze: its predictions for missing regions are likely influenced by memorized structural patterns. When the input structure diverges from these memorized patterns, reconstruction tends to fail. In contrast, the pretrained model is less dependent on such global pattern memory due to the use of CG representations, making it generalizable across diverse conformations. To extend the capability to reconstruct longer missing segments, we suppose that incorporating generative modeling frameworks, such as diffusion^40^ or flow matching^41^, should reduce reliance on memorized structural patterns and enable more robust structural sampling.

## Discussion

We developed StruCloze, a deep learning framework for reconstructing all-atom structures from CG models or partial biomolecular structures. StruCloze achieves strong reconstruction performance across various CG representations of proteins and nucleic acids, and can also generate reasonable predictions for entirely unresolved regions. Although targeted model finetuning improves performance for specific CG representations, the design of more representative CG schemes should also be regarded as a critical direction^42^.

Although adapting a general-purpose regime similar to AlphaFold3^11^, StruCloze differs significantly in both input and prediction objectives. Structure prediction models typically rely heavily on sequence-based features—such as multiple sequence alignments (MSAs) or embeddings from protein language models—which are essential to their success^43,44^. In contrast, our model primarily operates on low-resolution structural inputs and has minimal reliance on sequence information. Furthermore, traditional structure predictors aim to generate a single confident conformation for a given sequence, often resembling crystallographic structures. In contrast, StruCloze focuses on reconstructing the most plausible full-atom structure from any given partially resolved or coarse-grained input, even if the resulting structure may not be energetically favorable. This property may provide value for bridging static structures and ensemble modeling.

Numerous deep learning approaches have been proposed for backmapping CG models into full-atom representations. We propose that CG backmapping and missing region inpainting are inherently similar problems for deep learning model, both involving the completion of missing atomic details from partial structural data. By directly modeling atomic coordinates, a single neural network can generalize across these tasks. We demonstrate that deep learning models can be transferred between different CG representations and the two tasks with minimal finetuning and little loss in accuracy. This unified modeling framework and rapid adaptability make the method suitable for evolving CG representations, including newly designed CG schemes.

While StruCloze provides accurate all-atom estimations, it is important to note that these predictions do not capture structural dynamics; instead, they are more likely to resemble ensemble-averaged conformations, and repeated predictions yield nearly identical outputs. It means that prediction for locally dynamic biomolecules (e.g., disordered RNAs with extremely flexible nucleotides) may still be inaccurate. For side-chain dynamics in proteins, structure sampling tools such as AF2χ^27^ and DiffPack^45^ are more recommended, while sampling nucleotide conformations still rely on MD simulations.

StruCloze is currently implemented as a deterministic predictor, aiming for fast and accurate reconstruction. Incorporating this framework and its training strategies into generative models could unlock new possibilities in ensemble prediction and functional protein design. Predictive architecture has leaded to prediction towards only one conformation in inpainting missing regions, as have discussed in Results. Incorporating generative framework should reduce the reliance on memorization of certain structural patterns and make the model able to predict alternative conformations. Additionally, integrating richer protein and nucleic acid features may further improve model performance and extend its capabilities.

## Materials and Methods

### Dataset

We constructed a training dataset containing experimental biomolecular complexes for model pretraining and fine-tuning. The dataset includes protein monomers, multimers, nucleic acids and protein-nucleic acid complexes. We gathered all relevant PDB entries released before September 2021 and separated those containing nucleotides. Entries with missing or non-standard residues (and nucleotides) were removed. Protein entries were clustered at 40% sequence identity with MMseq2^46^, resulting in 36,014 structures, while all 16,912 structures containing nucleic acids were retained due to their limited availability. The final complex dataset comprises 52,570 structures.

For testing, we collected PDB entries released between 1 May 2022 and 12 January 2023. We removed systems containing over 2,048 tokens yielding 7,310 protein and 610 nucleic-acid-containing systems. Additionally, we included all-atom simulation data from Robustelli et al. for test on reconstructing protein trajectories.

The dataset compositions are summarized in Table 2.

**Table 2.** Training and test datasets.

| Stage | Entry type | Number of entries | Source |
| --- | --- | --- | --- |
| Train and validation (0.95:0.05) | Protein monomer | 25,352 | PDB<sup>47</sup> |
|  | Protein multimer | 10,504 |  |
|  | Nucleic acid | 3,902 |  |
|  | Complex | 12,812 |  |
|  | Total | 52,570 |  |
| Test | Protein | 7,310 |  |
|  | Nucleic acid | 78 |  |
|  | Complex | 678 |  |
|  | Total | 8,066 |  |
| External test (MD trajectories) | IDPs | 19 (*1000 conformations) | Robustelli et. al.<sup>36</sup> |
|  | ATLAS | 1,390 (*20 conformations) | Vander Meersche et al.<sup>37</sup> |
|  | Condensates | 3 (*1000 conformations) | Bülow et al.<sup>38</sup><br>Yasuda et al.<sup>22</sup> |

### Model Architecture

Conventional CG2AA algorithms—such as those reconstructing structures from dihedral angles or internal coordinates—often suffer from cumulative errors. Many recent approaches use the rigid-frame representation introduced by AlphaFold2, which, however, poses challenges when describing nucleotides. To overcome these issues, we developed StruCloze to directly predict Cartesian coordinates. StruCloze is adapted from the structure module of AlphaFold3 with several key differences:

1. **Diffusion Process:** We eliminated the diffusion process because our predictive task is considerably simpler than predicting structure from sequence alone. Iterative sampling would be computationally expensive with minimal performance gain.
2. **Sequence Representation:** Since our task benefits from a strong structural prior, the model does not heavily rely on complex sequence embeddings. Therefore, instead of using MSA information, we provide the model with one-hot encoded sequences.
3. **Feature Focus:** Atom-level features are prioritized over token-level features. Since structural priors already capture key token-level information, we reduced the number of token-level transformer layers to 4 (from 24 in AlphaFold3) and introduced additional atom-level transformers in both the encoder and decoder.
4. **Noisy Structures:** Unlike diffusion models that add scaled Gaussian noise to ground truth coordinates, our method perturbs reference coordinates (provided by CCD) based on coarse-grained representations, followed by local augmentations, including random rotations and translations. These perturbed coordinates serve as input noisy structures.

### Training Strategy

To ensure the transferability of our model across diverse coarse-grained representations, we designed a unified strategy for embedding CG information. Specifically, we first retrieved the reference atomic coordinates of each residue using its CCD code, and then aligned these coordinates by translating them according to the position of corresponding CG particle. For single-particle-per-residue representations (e.g., Cα/C4′ or residue center-of-mass), we applied random rigid-body transformations—both rotations and translations—to the shifted atomic coordinates for data augmentation. For multi-particle representations, we optimized the alignment between standard coordinates and CG particle positions by minimizing the angular deviation between corresponding inter-particle vectors, thereby preserving internal orientations.

This initialization scheme enables, in principle, a single pretrained model to reconstruct atomistic structures from a wide range of CG representations. However, in practice, the spatial separation between Cα/C4′ and the residue COM is often significant, making it challenging to achieve uniformly high accuracy with a single model. We therefore trained two specialized models in this study: StruCloze_CA and StruCloze_COM, each tailored to a specific CG representation.

Both models were trained through a three-stage pipeline (see Table 3). In the first stage, pretraining was performed using COM representations with an average translation perturbation of 1.5 Å for augmentation. This stage ran for 100 epochs on four NVIDIA 4090D GPUs, totaling approximately 76 hours. In the second stage, each model was fine-tuned on its target representation—Cα/C4′ or COM—using a reduced augmentation perturbation of 1.0 Å. Fine-tuning converged within 40 epochs, requiring about 30 hours. In the final stage, to enable inference on entirely unobserved residues (e.g., unresolved fragments in experimental structures), we introduced a masked self-supervised learning step based on the fine-tuned models. This stage, which required a longer convergence time, was trained for an additional 100 epochs. Notably, we observed that fine-tuned models converged as quickly as models trained from scratch, but consistently achieved lower final losses and superior structural fidelity.

**Table 3.** Parameter settings for three training stages.

| Parameters | Pretrain | Fine-tune | Masked Fine-tune |
| --- | --- | --- | --- |
| **Training profile** |  |  |  |
| Crop size (token) | 256 | 256 | 256 |
| Batch size | 16 | 16 | 16 |
| Epoch | 200 | 40 | 100 |
| **Loss profile** |  |  |  |
| Weight of MSE loss | 1/3 | 1/3 | 1/3 |
| Use LDDT loss | True | False | False |
| Use bond loss | False | True | True |
| **Data preprocess** |  |  |  |
| Ratio of masked residues(%) | 0.0 | 0.0 | 5.0 |
| Scale of random translation (Å) | 1.5 | 1.0 | 1.0 |

During training, we employed three cropping strategies for large biomolecular systems: spatial cropping, contiguous cropping on single chains, and multi-chain cropping, each generating fixed-size inputs of 256 residues. Spatial cropping, used in 40% of training iterations, randomly selected a central residue and retrieved its 256 nearest neighbors. Single-chain cropping randomly selected a single chain from a complex and extracted a continuous segment. In multi-chain cropping, we sampled segments from multiple chains within an entry, jointly yielding 256 residues. The latter two strategies were each used in 30% of training iterations.

### Evaluation Metrics

A primary metric for evaluating the reconstruction of all-atom structures is the root mean squared deviation (RMSD) of Cartesian coordinates, defined as:

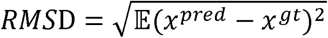

where *x^pred^* and *x^gt^* denotes atom coordinates of predicted and ground truth structures respectively. We calculated RMSD on all heavy atoms to evaluate the overall reconstruction accuracy.

RMSD represents global reconstruction quality well but is often relatively insensitive to local errors such as steric overlaps. Therefore, we calculated clash ratio to evaluate local geometric plausibility between non-bonded neighboring atom pairs. Typically, steric clash is defined as two atoms being closer than the sum of their Van Der Waals (VDW) radii^48^. We define the clash ratio *r_c_* as the fraction of atom pairs that exhibit steric clashes among those that are 3–8□Å apart in the ground-truth structure:

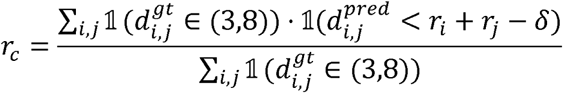

In which 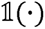 is an indicator function equaling 1 if condition is true, 0 otherwise. 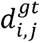 and 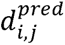 denotes distance between atom i and j in ground truth and predicted structures separately. *r_i_*, *r_j_* are VDW radii of atoms i and j, while *δ* = 0.4 is an allowed minor overlap.

Besides distances, orientations of backbones and side chains also account for the quality of biomolecular structures. To account for this part, we calculated accuracy for protein χ1 and χ2, in which prediction is considered correct if the deviation from the experimental angle is less than 30°. For nucleic acids, we calculated the pseudo angles χ(p1) to χ(p5) for the ribose backbone (Table 4), the sugar pucker (whether nucleotide adopts C2’-endo or C3’-endo conformation) is defined as follows:

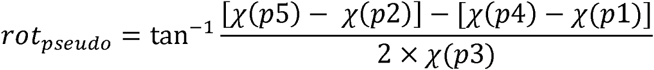

where C3’-endo conformations correspond to *rot_pseudo_* near 0° while C2’-endo typically lies near 180°^49^.

**Table 4.** Definition of pseudo angles in nucleic acids.

| Pseudo angle | Atom 1 | Atom2 | Atom3 | Atom4 |
| --- | --- | --- | --- | --- |
| $\chi(p1)$ | O4' | C1' | C2' | C3' |
| $\chi(p2)$ | C1' | C2' | C3' | C4' |
| $\chi(p3)$ | C2' | C3' | C4' | O4' |
| $\chi(p4)$ | C3' | C4' | O4' | C1' |
| $\chi(p5)$ | C4' | O4' | C1' | C2' |

To evaluate base pairing, we assessed the quality of predicted hydrogen bonds using precision and recall. Given the large imbalance between true negatives (TN) and true positives (TP), overall accuracy is always close to 1 and thus uninformative; we therefore did not report accuracy in this context.

## Acknowledgement

This work was supported by Shanghai Municipal Science and Technology Major Project, partially by SJTU Kunpeng & Ascend Center of Excellence, the Center for HPC at Shanghai Jiao Tong University, and the National Key Research and Development Program of China (2025YFA0921001 and 2023YFF1205102), the Fundamental Research Funds for the Central Universities (YG2023LC03), and the National Natural Science Foundation of China (32171242).

## Data and Code Availability

Structure data used in model training and testing are available at https://doi.org/10.5281/zenodo.15524132. Source code and model parameter files are available at https://github.com/Junjie-Zhu/StruCloze.

## Supplementary figures

**Figure S1.**
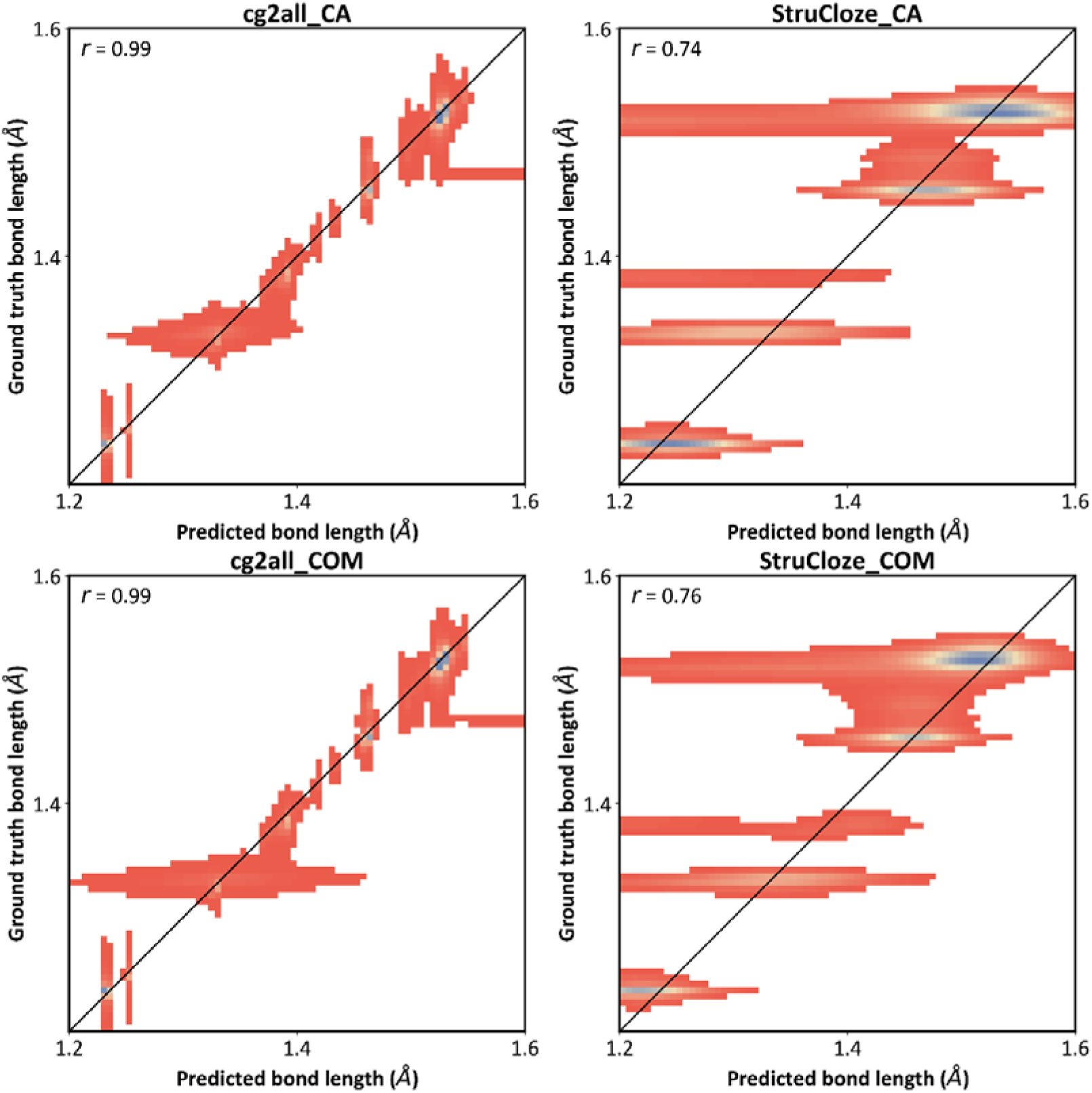
Bond distribution of structures reconstructed by StruCloze and cg2all.

**Figure S2.**
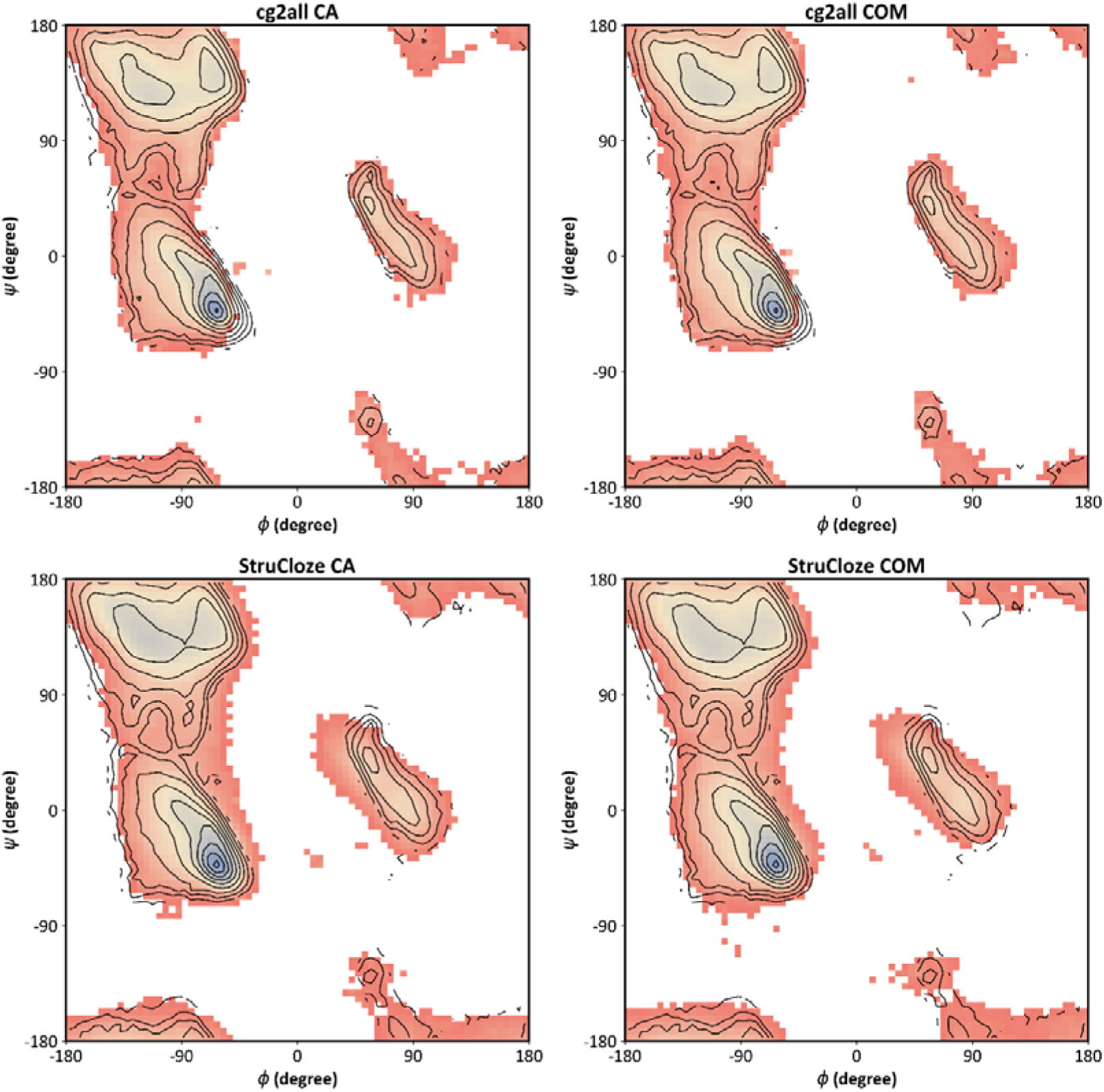
Ramachandran plots for cg2all and StruCloze predicted protein structures. Distribution for experimental structures are shown as black contours.

**Figure S3.**
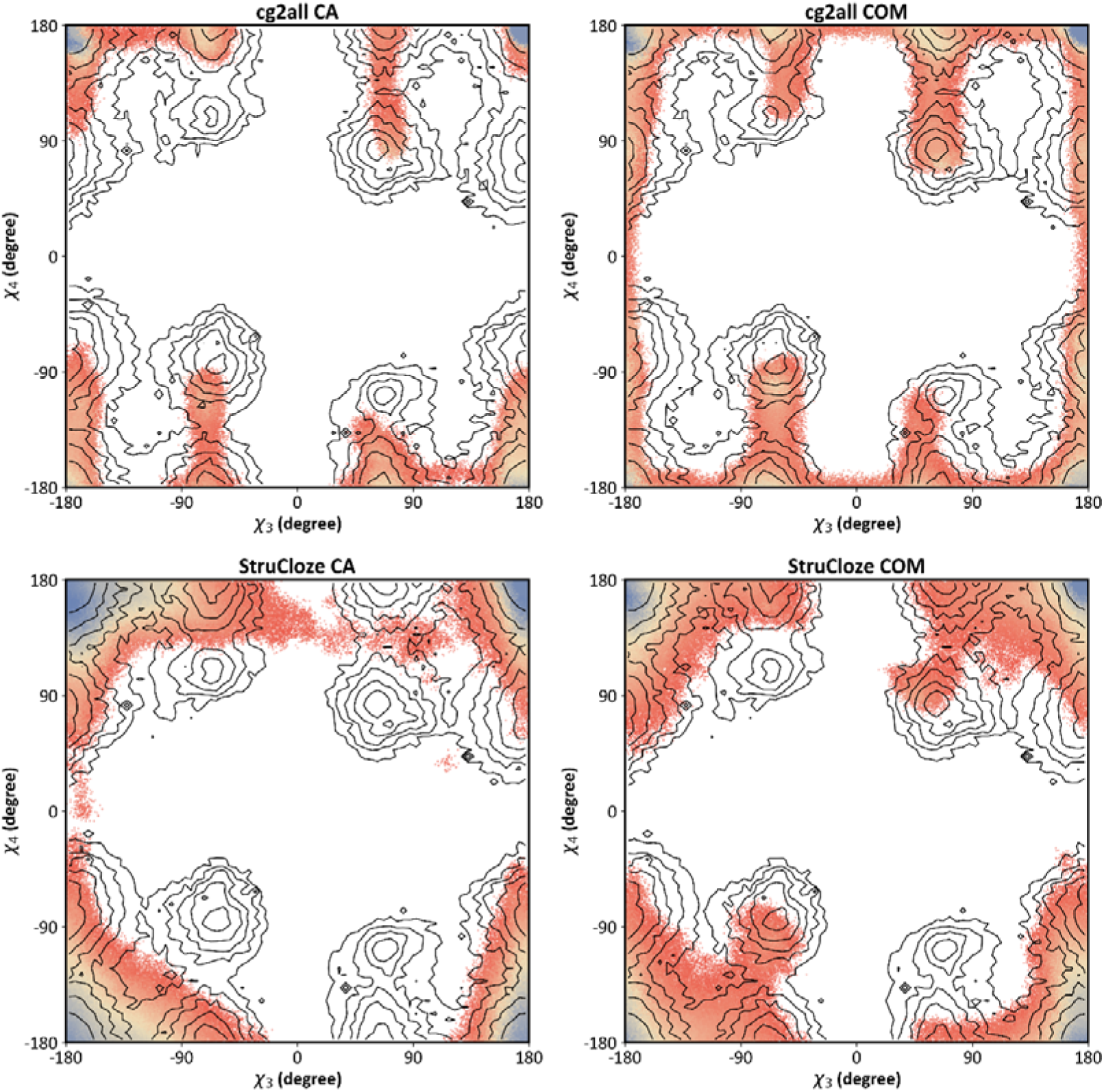
χ3-χ4 distributions for cg2all and StruCloze predicted protein structures. Distribution for experimental structures are shown as black contours.

**Figure S4.**
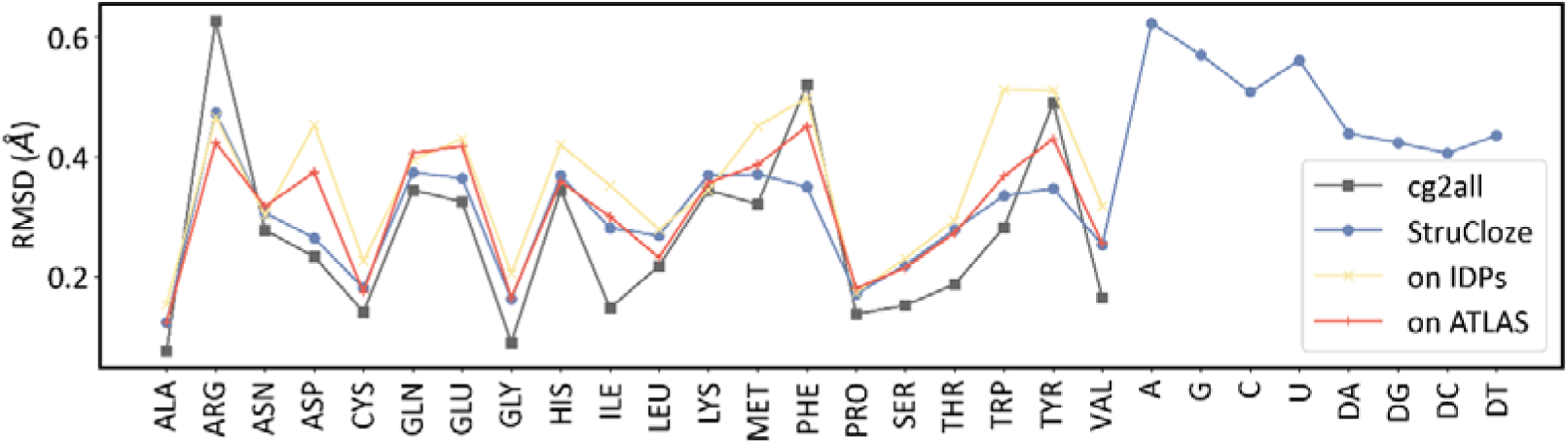
Residue-level reconstruction RMSD.

**Figure S5.**
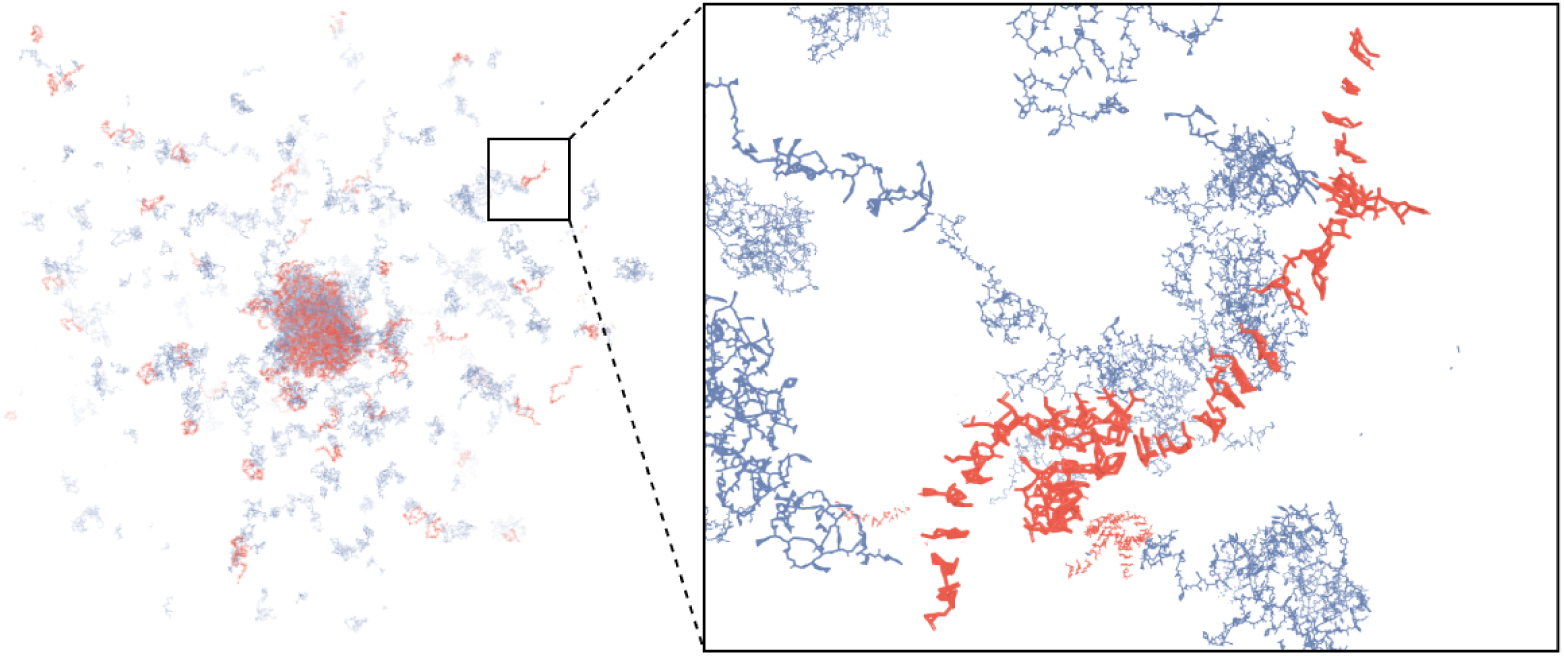
Reconstructed structure for one snapshot in simulation of FUSRGG3 and polyU40. Local structure of RNA components are hard to predict.

